# Cryogenically preserved RBCs support gametocytogenesis of *Plasmodium falciparum* in *vitro* and gametogenesis in mosquitoes

**DOI:** 10.1101/405001

**Authors:** Ashutosh K. Pathak, Justine C. Shiau, Matthew B. Thomas, Courtney Murdock

## Abstract

**Background:** The malaria Eradication Research Agenda (malERA) has identified human-to-mosquito transmission of *Plasmodium falciparum* as a major target for eradication. The cornerstone for identifying and evaluating transmission in the laboratory is small membrane feeding assays (SMFAs) where mature gametocytes of *P. falciparum* generated *in vitro* are offered to mosquitoes as part of a blood-meal. However, propagation of “infectious” gametocytes requires 10-12 days with considerable physico-chemical demands imposed on host RBCs and thus, “fresh” RBCs that are ≤1-week old post-collection are generally recommended. However, in addition to the costs, physico-chemical characteristics unique to RBC donors may confound reproducibility and interpretation of SMFAs. Cryogenic storage of RBCs (cryo-preserved RBCs herein) is approved by the European and US FDAs as an alternative to refrigeration (4°C) for preserving RBC quality and while cryo-preserved RBCs have been used for *in vitro* cultures of other *Plasmodia* and the asexual stages of *P. falciparum*, none of the studies required RBCs to support parasite development for >4 days.

**Results:** Using the standard laboratory strain, *P. falciparum* NF54, we first demonstrate that cryo-preserved RBCs preserved in the gaseous phase of liquid nitrogen and thawed after storage for 1, 4, 8 and 12 weeks, supported gametocytogenesis *in vitro* and subsequent gametogenesis in *Anopheles stephensi* mosquitoes. Using data from 11 SMFAs and RBCs from 4 separate donors with 3 donors re-tested following various periods of cryo-preservation, we show that overall levels of sporogony in the mosquito, as measured by oocyst prevalence and burdens in the midguts and sporozoites in salivary glands, were similar or better than using ≤1-week old refrigerated RBCs. Additionally, the potential for cryo-preserved RBCs to serve as a universal substrate for SMFAs is shown for a Cambodian isolate of *P. falciparum*.

**Conclusions:** Considering the suitability of cryo-preserved RBCs for *P. falciparum* SMFAs, we suggest guidelines for their use and how they can be integrated into an existing laboratory/insectary framework with the potential to significantly reduce running costs and provide greater reliability. Finally, we discuss scenarios where cryo-preserved RBCs may be especially useful in enhancing our understanding and/or providing novel insights into the patterns and process underlying human-to-mosquito transmission.

## Background

Incidence of malaria due to *Plasmodium falciparum* has seen a steady decline in the last 15-20 years, enabled primarily by the synergy between drug-combination therapies for case/disease management and vector control programs [1, 2]. Since transmission to the mosquito is thought to be a severe bottleneck in the parasite’s life history, considerable effort has been channeled towards identifying parasite and vector traits influencing transmission to the mosquito with the aim of designing and testing small molecule inhibitors and vaccines [3-6]. The experimental model for evaluating basic parasite life history or the efficacy of various interventions typically involve small-membrane feeding assays (SMFAs), where mature male and female gametocytes of *P. falciparum* cultured *in vitro* are supplemented with naïve RBCs and offered as part of a blood-meal to female *Anopheles* spp. mosquitoes. These assays are then followed by the collection of various measures of parasite fitness in the mosquito vector [7-11]. However, in the 40 years since Trager and Jensen’s first description of the methodology, *in vitro* cultures of transmission-competent parasite stages (the definitive measure of parasite fitness) have proven to be an arduous and labor-intensive venture with the precise mechanism(s) regulating the induction of gametocytogenesis remaining largely elusive [12-15]. Currently available methodology is a synthesis of several tools and techniques that have been elegantly summarized in two recent publications [13, 15].

Relative to other etiological agents of human malaria, *P. falciparum* is unique in its life-history in blood wherein the asexual replication period lasts 2 days during which approximately 15% of the resulting parasite progeny is pre-destined to invade an RBC and enter an irreversible path of sexual differentiation lasting an additional 10-12 days before maturation into male and female gametocytes capable of infecting a mosquito. Most references outlining methods for growing sexual stages in RBCs *in vitro* emphasize the need for maintaining the sexual stages in fresh RBCs (i.e., storage under refrigeration for no more than 1-week from the day of collection), as the RBC must be able to support gametocytogenesis both physically and energetically during the 10 to 12-day differentiation period [13, 15-18]. However, ensuring a regular supply of commercially sourced RBCs can quickly prove to be prohibitively expensive. Additionally, variation introduced by differences in storage time of RBCs and across blood donors can potentially carry significant consequences for the reproducibility of downstream experiments and across laboratories for validating vaccine and drug efficacy for instance [9, 19-21].

Despite >100 years of experience in bio-banking, identifying the true “shelf-life” of refrigerated RBCs is controversial even in case of human blood transfusions [22]. While it may be possible to restore some aspects of RBC metabolism upon rejuvenation of packed RBCs with the addition of fresh nutrient-containing media, most of the storage-associated changes are simply irreversible [23-25]. As an alternative, cryo-preservation of RBCs in glycerol-based cryo-protectants (storage at −65°C or below) has been suggested using procedures approved by both the United States Food and Drug Administration and the European Council [26, 27].

While cryo-preserved RBCs had been shown to support the growth of other etiological agents of human malaria such as *P. vivax*, a recent study demonstrated that growth rates of the asexual blood stages of *P. falciparum* in cryogenically-preserved RBCs (−196°C in the gaseous phase of liquid nitrogen) was almost identical to freshly collected blood *in vitro* [28, 29]. However, none of these studies required the RBCs to sustain parasite growth for more than four days. According to current FDA guidelines, cryo-preserved RBCs thawed under sterile conditions can retain physicochemical properties for up to (at least) 14 days following the thaw date [26]. Considering how this timeframe encompasses the duration required for gametocytogenesis of *P. falciparum*, we hypothesized that cryo-preserved RBCs should also provide a suitable substrate for the culture of sexual stages of *P. falciparum*. In the current study, we were interested in determining if cryo-preserved RBCs support gametocytogenesis, and if the resulting mature gametocytes undergo gametogenesis using SMFAs with *Anopheles stephensi* as the definitive measure of fitness. We also addressed whether infectiousness was retained following extended periods of cryo-preservation. Lastly, although we accomplished these objectives primarily with the standard lab strain (NF54) of *P. falciparum*, we also tested the potential of cryo-preserved RBCs to serve as a substrate for supporting growth and differentiation of infectious stages of a Cambodian isolate of *P. falciparum*, CB132 [30].

## Results

### Gametocytogenesis of *P. falciparum in vitro*

Mature gametocytemia *in vitro* generally increased over time with detectable levels achieved by 12 days post-infection (figure 1). Overall trends were independent of storage method (Odds ratios (OR)= 1.31, standard error (se)=0.41, *p*=0.4) or duration (OR=1.01, se=0.04, *p*=0.72) with most of the random variation is explained by differences between experimental block and negligible contribution by RBC donor (Table 1).

**Table 1:** outputs of the statistical models

| Dependent variable → | Gametocytemia *in vitro* (male + female stage V) |  |  | Oocyst prevalence in mosquito midguts |  |  | Sporozoite prevalence in salivary glands |  |  | Oocyst abundance (infected and uninfected midguts) |  |  | Oocyst intensity (infected midguts only) |  |  |
| --- | --- | --- | --- | --- | --- | --- | --- | --- | --- | --- | --- | --- | --- | --- | --- |
| Summary statistics → | IRR<sup>a</sup> | std. Error | *p-value* | OR<sup>a</sup> | std. Error | *p-value* | OR<sup>a</sup> | std. Error | *p-value* | IRR<sup>a</sup> | std. Error | *p-value* | IRR<sup>a</sup> | std. Error | *p-value* |
| Fixed Effects |  |  |  |  |  |  |  |  |  |  |  |  |  |  |  |
| (Intercept) | 0.00 | 0.00 | <.001 | 1.09 | 0.52 | 0.86 | 0.82 | 0.50 | 0.75 | 1.96 | 0.85 | 0.43 | 5.19 | 0.48 | <.001 |
| 4°C RBCs vs Cryo-RBCs | 1.31 | 0.41 | 0.40 | 1.05 | 0.38 | 0.89 | 1.35 | 0.84 | 0.63 | 1.20 | 0.71 | 0.80 | 1.12 | 0.36 | 0.75 |
| Duration of storage | 1.01 | 0.04 | 0.72 | 0.89 | 0.06 | 0.07 | 0.93 | 0.08 | 0.36 | 0.86 | 0.10 | 0.13 | 0.95 | 0.07 | 0.48 |
| Experimental design |  |  |  |  |  |  |  |  |  |  |  |  |  |  |  |
| Total flasks/mosquito infections (SMFAs), nested within experimental block | 11 |  |  | 11 |  |  | 10<sup>b</sup> |  |  | 11 |  |  | 11 |  |  |
| Total number of experimental blocks | 5 |  |  | 5 |  |  | 5 |  |  | 5 |  |  | 5 |  |  |
| Total no. of RBC donors tested | 4 |  |  | 4 |  |  | 4 |  |  | 4 |  |  | 4 |  |  |
| Total number of observations | 51 |  |  | 283 |  |  | 235 |  |  | 283 |  |  | 157 |  |  |
| Random effects |  |  |  |  |  |  |  |  |  |  |  |  |  |  |  |
| <u>Group structure and variance for random effects</u> |  |  |  |  |  |  |  |  |  |  |  |  |  |  |  |
| Flasks/SMFAs nested within experimental block | 0.00 |  |  | 0.00 |  |  | 0.26 |  |  | 0.43 |  |  | 0.10 |  |  |
| Experimental block | 0.16 |  |  | 0.35 |  |  | 0.00 |  |  | 0.77 |  |  | 0.40 |  |  |
| RBC donor | 0.00 |  |  | 0.17 |  |  | 0.02 |  |  | 0.46 |  |  | 0.21 |  |  |
| <u>Overdispersion tests</u> |  |  |  |  |  |  |  |  |  |  |  |  |  |  |  |
| Dispersion ratio | 0.98 |  |  | 1.23 |  |  | 1.00 |  |  | 0.75 |  |  | 1.07 |  |  |
| p-value | 0.50 |  |  | 0.29 |  |  | 0.40 |  |  | 1.00 |  |  | 0.27 |  |  |
| Notes |  |  |  |  |  |  |  |  |  |  |  |  |  |  |  |
| GLMM family (link) | poisson (log) |  |  | binomial (logit) |  |  | binomial (logit) |  |  | negative binomial (log) |  |  | negative binomial (log) |  |  |
| Abbreviations<sup>a</sup> | IRR= Incidence rate ratios, NA=not applicable, OR=Odds ratios |  |  |  |  |  |  |  |  |  |  |  |  |  |  |
| Comments<sup>b</sup> | <sup>b</sup>no of infections is lower as sporozoite prevalence was not available from 1 infection |  |  |  |  |  |  |  |  |  |  |  |  |  |  |

**Figure 1:**
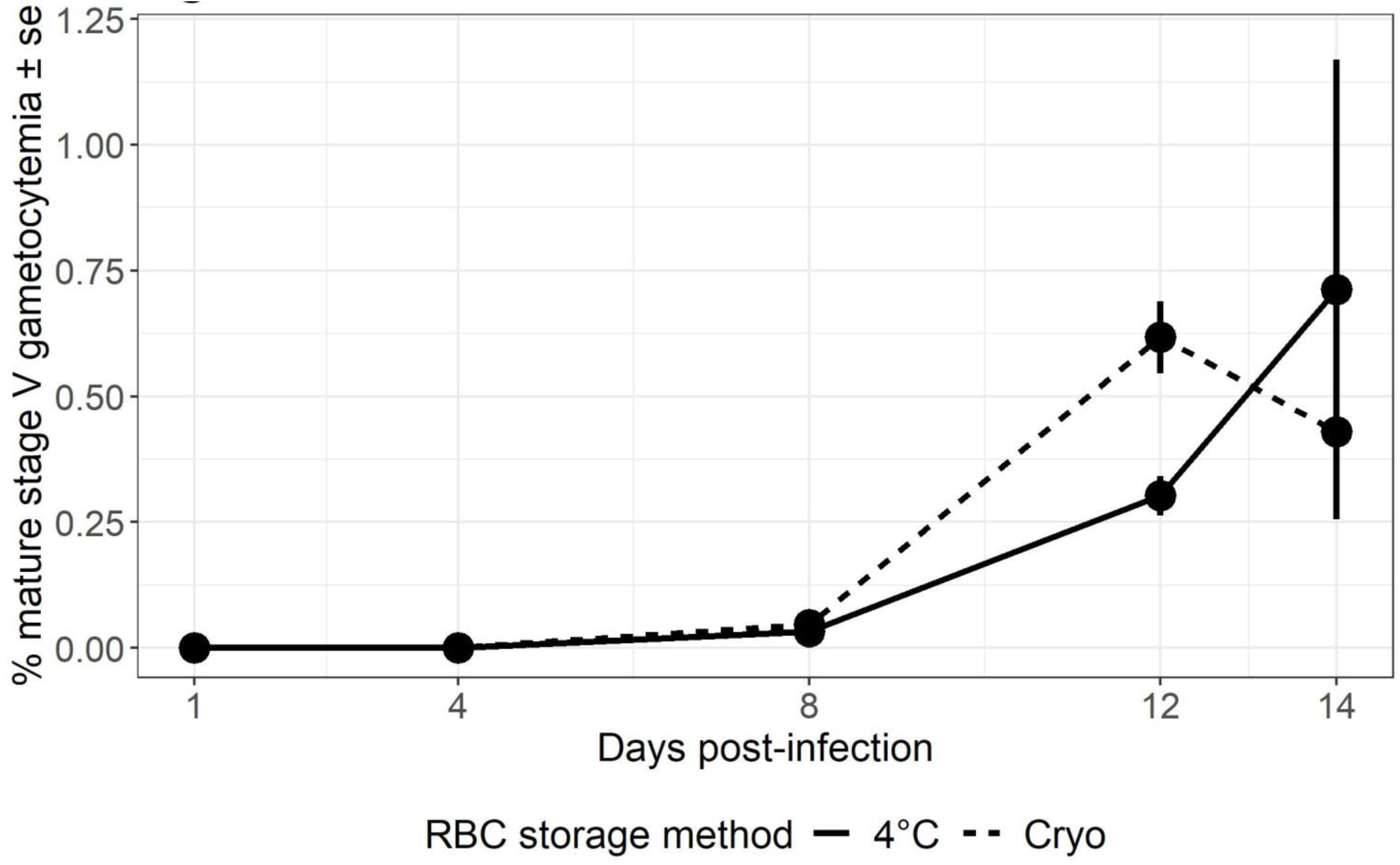
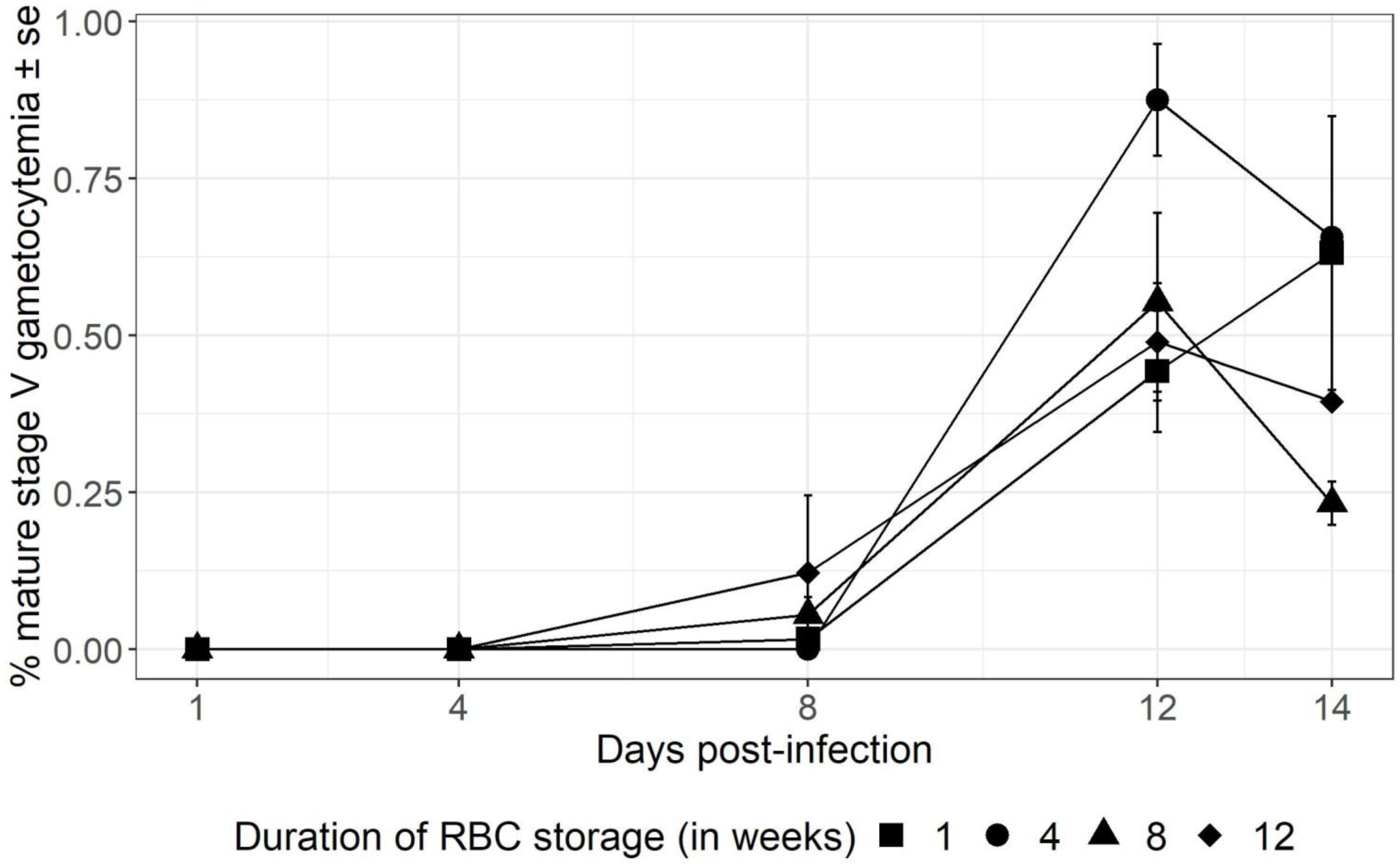
a) Rates of mature gametocytemia of *P. falciparum* NF54 cultured *in vitro* in refrigerated (continuous line) or cryo-preserved (dashed line) RBCs and b), rates of gametocytemia in RBCs cryo-preserved within 3-4 days of collection from each donor and thawed after 1 (closed squares), 4 (closed circles), 8 (closed triangles) and 12 weeks (closed diamonds) of storage. Data represents mean ± standard errors (se).

### Prevalence of *P. falciparum* in midguts of *An. stephensi* mosquitoes

The mean (±se) prevalence of mosquitoes infected with *P. falciparum* oocysts (NF54) was 53±6.4% (n=11 SMFAs, mean mosquito midguts sampled/SMFA= 26 (range= 17-44), total number of mosquito midguts sampled=283) (figure. 2a). While prevalence was independent of storage method (OR=1.05, se=0.38, *p*=0.89), the ability of cryo-preserved RBCs to culture infectious gametocytes showed a marginally insignificant decline with duration of storage (OR=0.89, se=0.18, *p*=0.07) with much of the unexplained variation represented by differences across experimental blocks, although some could also be attributed to RBC donor characteristics (Table 1 and additional file 3).

**Figure 2:**
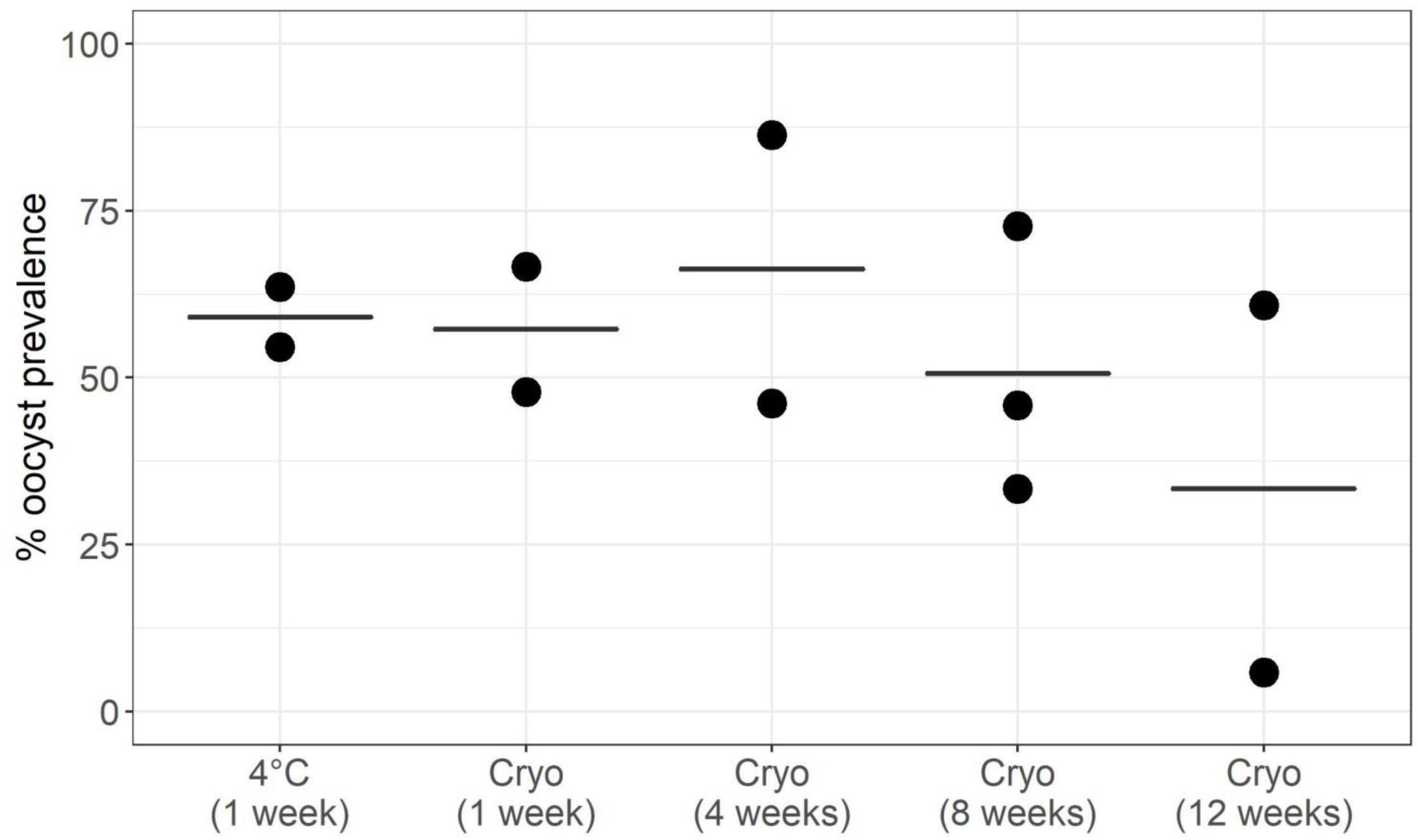
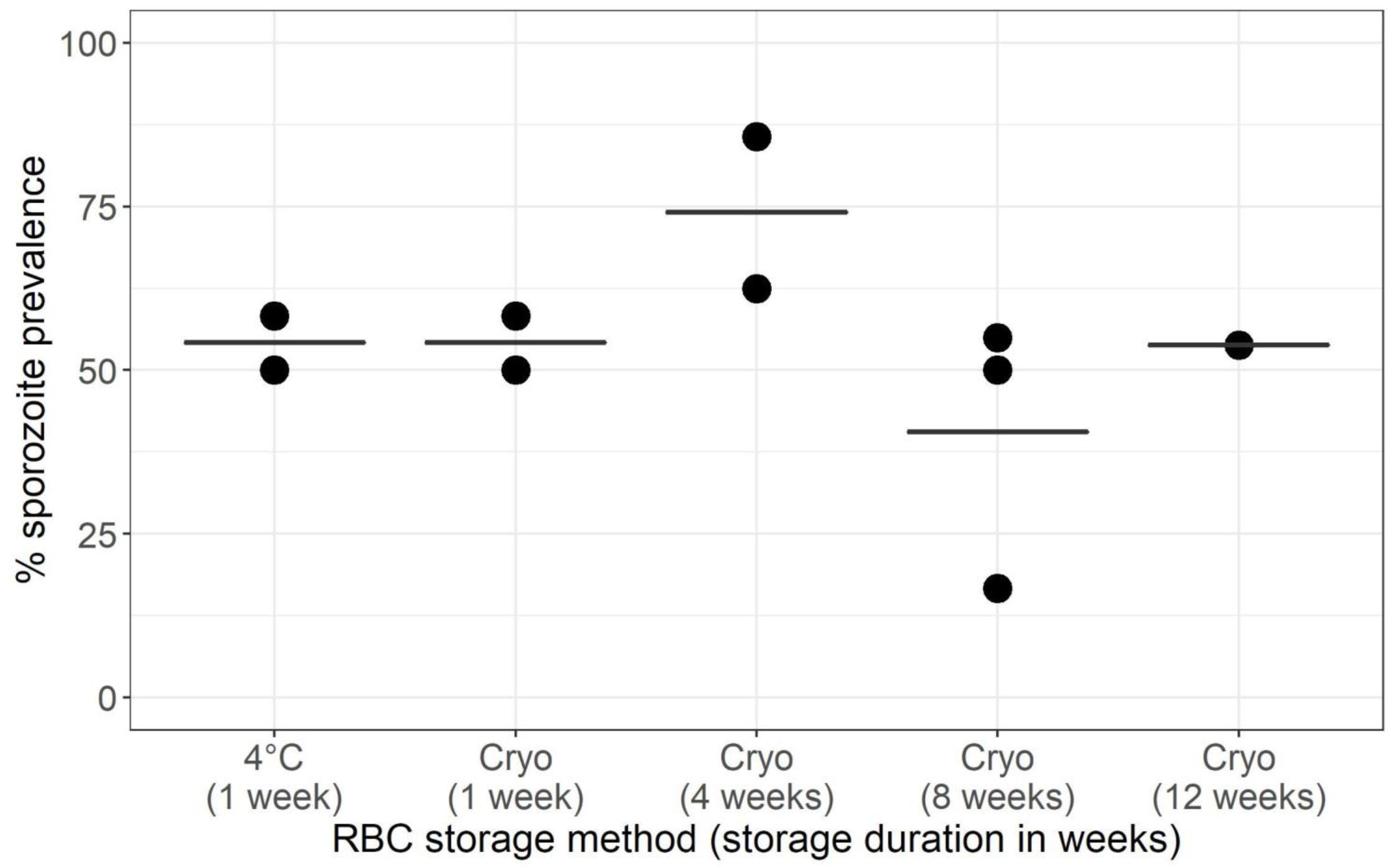
Prevalence of a) oocysts and b) sporozoites in midguts and salivary glands respectively of *An. stephensi* mosquitoes infected with ∼0.1% mature gametocytemia of *P. falciparum* NF54. Each data point represents an SMFA where mosquitoes were provided a blood-meal spiked with mature ex-flagellation competent gametocytes collected from the flasks depicted in Figure 1 and cultured in RBCs refrigerated (4°C), cryo-preserved (“Cryo”) for ≤1 week, or following storage for 4, 8 and 12 weeks. See additional file 2 for graphical representation of the study design and additional file 4 demonstrating variation in oocyst prevalence between experimental blocks.

### Prevalence of *P. falciparum* in the salivary glands of mosquitoes

The mean (±se) prevalence of mosquitoes carrying sporozoites of *P. falciparum* NF54 in the salivary glands was 54±5.3% (n=10 SMFAs, mean salivary glands sampled/SMFA= 24 (range= 20-30), total number of salivary glands sampled=235) (figure 2b). Overall trends in sporozoite prevalence were independent of storage method (OR=1.35, se=0.84, p=0.63) or duration (OR=0.93, se=0.08, p=0.36) (Table 1). Unlike the midguts however, most of the random variation was caused by differences among infections with negligible contributions from experimental block or RBC donor (Table 1).

### Oocyst abundance and intensity in the midguts of mosquitoes

Mean (±se) oocyst abundance (oocyst counts from all sampled midgut) was 5.4±2.53 (n=11 SMFAs, mean mosquito midguts sampled/SMFA= 26 (range= 17-44), total number of mosquito midguts sampled=283) while oocyst intensity (oocyst counts from infected midguts only) was 7.75±2.85 (n=11 SMFAs, range of infected midguts/SMFA= 1-28, total number of mosquito midguts sampled=157) (figure 3). While both abundance and intensity were independent of storage method (Incidence rates ratio for abundance (IRR)= 1.20, se=0.71, p=0. 80, IRR for intensity=1.12, se=0.36, p=0.75) or duration (IRR for abundance= 0.86, se=0.1, p=0.13, IRR for intensity=0.95, se=0.07, p=0.48) (Table 1), most of the random variation was generated by the differences between experimental blocks with some contribution from RBC donor-specific characteristics and the least from the variation among infections.

**Figure 3:**
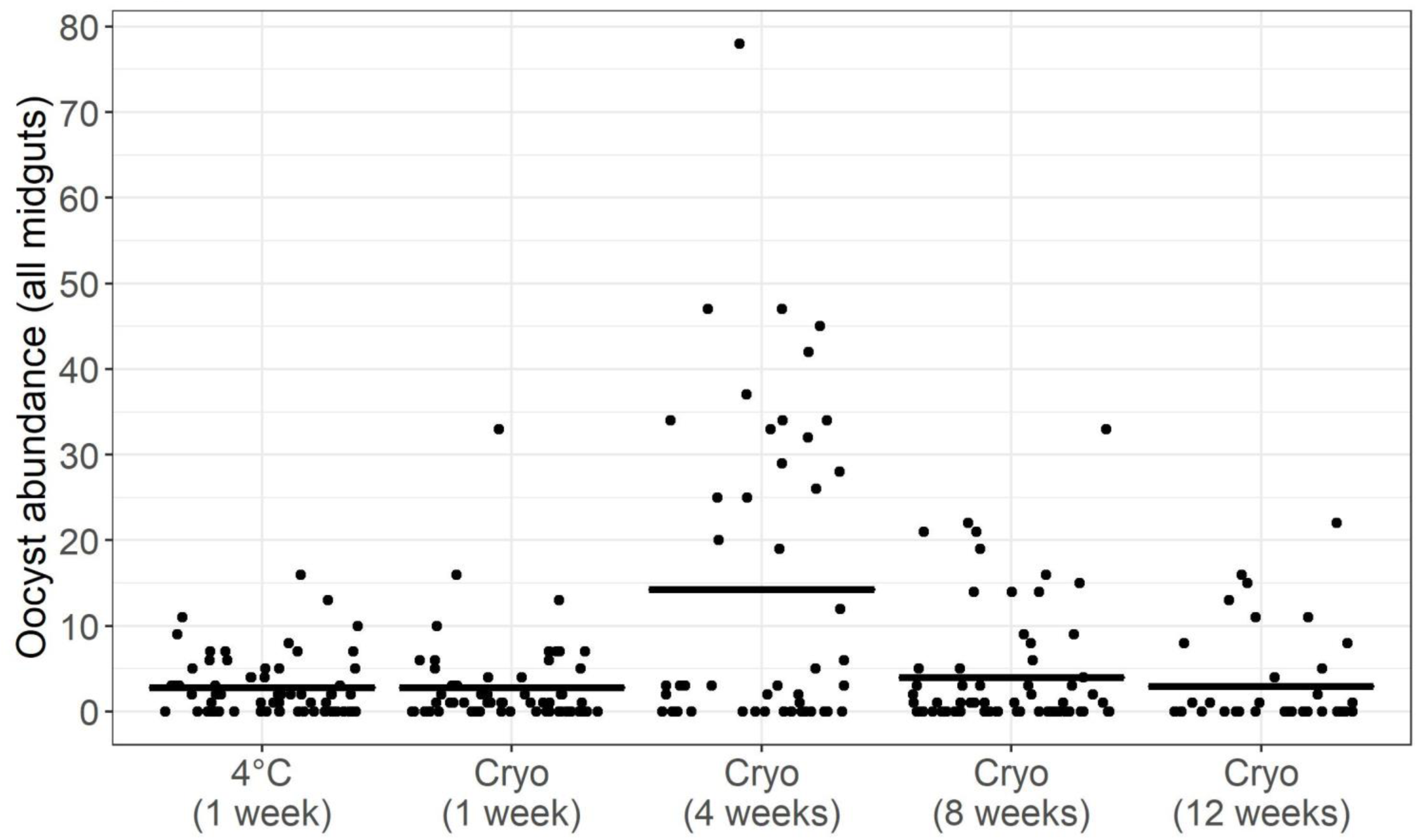
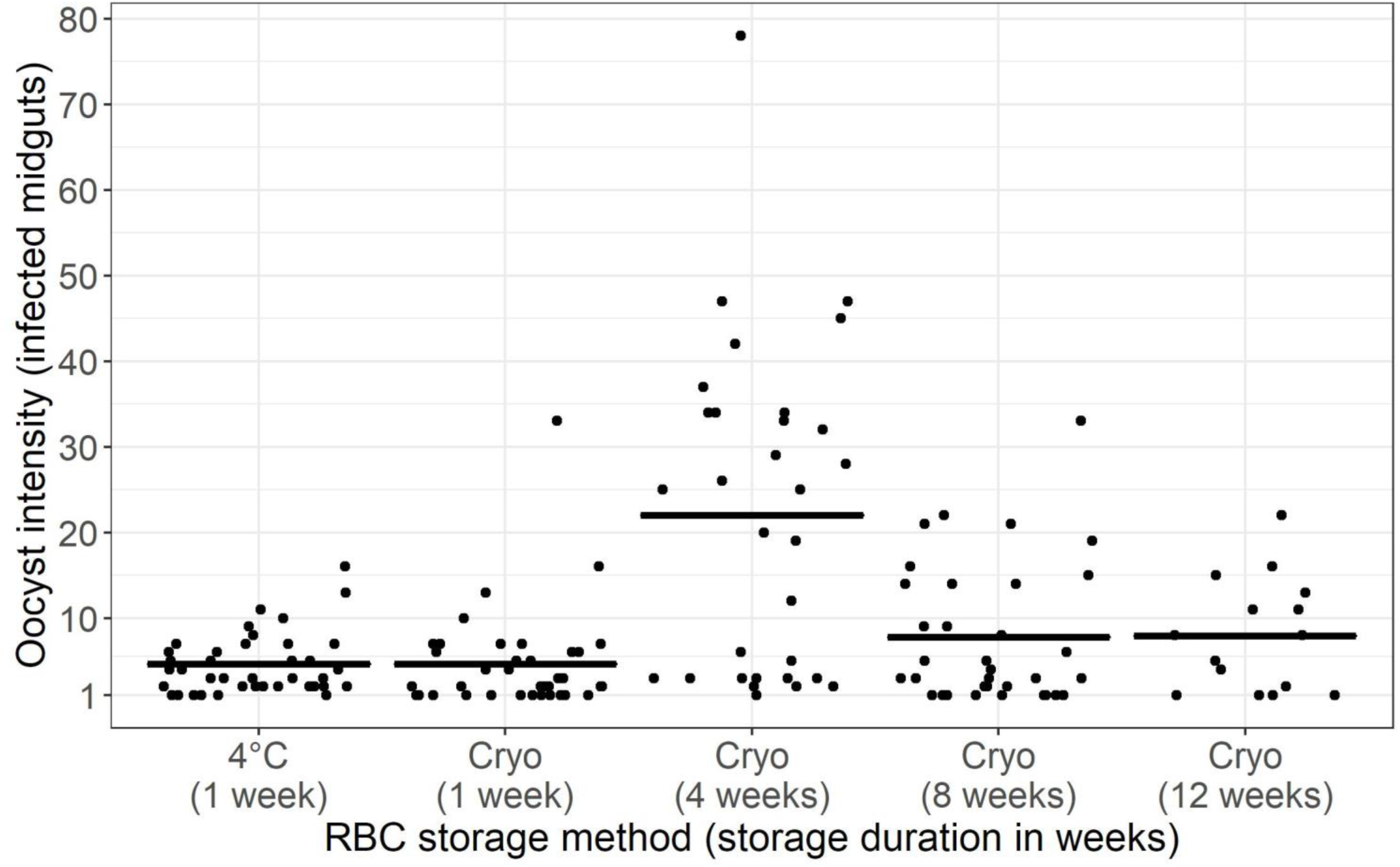
Oocyst a) abundance (oocyst counts from all midguts regardless of infection status) and b) intensity (oocyst counts from infected midguts only) in mosquitoes depicted in figure 2a. Horizontal bars represent group means with each data point representing oocyst counts from an individual mosquito midgut. For visualization purposes, counts were jittered horizontally to 40% but not vertically to maintain alignment with the gradient on the y-axis.

### Gametogenesis of Cambodian isolate *P. falciparum* (CB132)

To determine if cryo-preservation can support infectious cultures of other strains of *P. falciparum,* RBC donor number 4 (additional file 2b) used for the NF54 cultures above was used as a substrate for sexual stage cultures of a clinical isolate of Cambodian origin (*P. falciparum* CB132 [30]) following storage periods of 6 (n=35 midguts and 23 salivary glands) and 8 weeks (n=49 midguts) (additional file 5). Statistical analyses were not performed due to the lack of replication in the study design.

## Discussion

Although cryogenically preserved RBCs have been shown to be suitable for culturing other *Plasmodium* species, including the asexual stages of *P. falciparum*, none of the studies ran *P. falciparum* cultures for more than four days [28, 29]. In the current study, we demonstrated that in addition to supporting proliferation of the asexual stages for the first 4 days, cryo-preserved RBCs were able to maintain their “qualities” as a substrate for the additional 10-12 days required for progression through the various stages of gametocytogenesis. Crucially, the fitness of the resulting sexually mature gametocytes was confirmed by successful gametogenesis and sporogony in *An. stephensi* mosquitoes.

Neither cryo-preservation nor duration of storage up to 12 weeks affected the ability of cryo-preserved RBCs to provide a substrate for parasite growth and differentiation of asexual stages of *P. falciparum* NF54 into mature gametocytes *in vitro*. Although rates of gametocytemia were apparently higher in cryo-preserved RBCs with a decline in concentrations following a peak at 12 days post-infection (figure 1), this non-linearity was accounted for by the random variation between experimental blocks (Table 1). The most obvious difference between the blocks was the growth rates of asexual stages in the asexual feeder flasks used to initiate the gametocyte cultures in each experimental block. Although statistical models suggested a positive association with the proportions of RBCs infected with parasites at the late trophozoite stage and subsequent rates of gametocytemia, our study design precluded more robust analyses of this relationship. Indeed, future experiments with synchronized asexual feeder cultures seeded at various starting densities should help tease apart the underlying relationships, manipulation of which could help design more streamlined regimes for inducing gametocytogenesis. Nevertheless, the relationship between late trophozoites in asexual feeder flasks and future gametocytemia has been suggested by others [13] and taken together, our observations lend further support to the importance of considering parasite fitness at the asexual stages and subsequent gametocytogenesis *in vitro*.

The rates of gametocytemia *in vitro* were not entirely reflected in the patterns of oocyst prevalence of *P. falciparum* NF54 in midguts of *An. stephensi* mosquitoes. Although not significant, the ability of cryo-preserved RBCs to support infectious gametocytes suggested a gradual decline after 12 weeks of storage. Although this trend was likely driven by one of the two biological replicates representing 12 weeks of cryo-preservation, to be conservative, we suggest restricting the use of cryo-preserved RBCs to 8-weeks if the objective is to perform SMFAs. Whether the observed low prevalence is a donor-specific characteristic would have to be confirmed with more RBC donors representing 12 weeks of cryo-preservation, however, when considered along with the other 2 infections performed within the same experimental block (additional file 3), mean oocyst prevalence was lower relative to the other experimental blocks. Indeed, refitting statistical models after excluding this infection resulted in a significant reduction in random RBC donor-introduced variation (0.17 in Table 1 to 0.04, remaining output not shown) but less so for the experimental block (0.35 to 0.19), suggesting that in general, while RBC donor characteristics may not be a significant source of variability in oocyst prevalence, a blocked experimental design may warrant careful consideration for SMFAs to aid robust interpretation of the results. Future experiments should help tease apart if this variability between experimental blocks is the result of 1) subtle differences between mosquito generations and/or 2) final parasite density and/or infectiousness which can vary significantly between generations [9]. Sporozoite prevalence in the salivary glands is independent of storage method and duration and largely mirrors the profiles of oocyst prevalence in the midguts, albeit with negligible random variation contributed by experimental blocks or RBC donor, with the majority resulting from differences between the 10 infections *per se*. However, this variability was being driven primarily by 1 outlier infection where prevalence was low for one of the RBC donors tested following 8 weeks of cryo-preservation, the exclusion of which was able to account for most if not all the previously unexplained variability between infections after refitting the model (data not shown). Similarly, neither oocyst abundance nor intensity were affected by maturation of gametocytes in 1-week old refrigerated or 1 to 12-week old cryo-preserved RBCs, with significant random variation across experimental blocks, infections and RBC donor. It is important to note that although we follow parasitological convention in differentiating between oocyst abundance and intensity [31], “intensity” has also been used to describe abundance in other studies [7, 9].

Lastly, preliminary experiments suggest that cryo-preserved RBCs should, with further refinements, support differentiation of other strains of *P. falciparum*. 6- and 8-week old cryo-preserved RBCs from one of the donors previously verified to support gametocytogenesis of NF54 was used to generate transmission-competent gametocytes of a Cambodian isolate of *P. falciparum*, CB132 (additional file 5) [30]. The lower levels of infection relative to the standard laboratory isolate, NF54, could reflect differences in “infectiousness” between the two strains while the reduction in oocyst prevalence at 8 weeks compared to 6 weeks could be the result of random variation between experimental blocks as seen for NF54.

## Conclusions

Gametocytogenesis of *P. falciparum* in RBCs cryo-preserved within 1 week of collection retained infectivity to mosquitoes for up to 12 weeks of storage, at levels similar or better than the “gold-standard” of using refrigerated 1-week old RBCs, with the observed patterns likely being governed by similar rate-limiting factors as refrigerated RBCs. However, while our results suggest that cryo-preservation might be suitable for culturing mature gametocytes *in vitro* for up to 12 weeks, their infectiousness to mosquitoes may decline after 8 weeks for some RBC donors. From an empirical perspective, the ability to culture parasites in the same donor of RBCs will increase the reproducibility of SMFAs, especially in scenarios where the contribution of RBC donor may warrant consideration. Further, the use of cryo-preserved RBCs for SMFAs is not restricted to that of the canonical NF54 isolate of *P. falciparum* and should also be applicable to other strains of *P. falciparum*, as we demonstrated for a Cambodian isolate, CB132 (additional file 6). From an economic perspective, 8 weeks offers a significant improvement over the current practice of using RBCs that are ≤1-week old, with the potential to therefore reduce the costs by almost 7-fold. Moreover, considering how this period also aligns with the 2-month duration suggested for storing blood for maintaining breeder colonies of mosquitoes ([32] and unpublished observations), it is conceivable for laboratories to integrate SMFAs using cryo-preserved RBCs into the general mosquito rearing schedule. For instance, we routinely aliquot RBCs for cryo-preservation within 3-4 days of collection for culturing parasites *in vitro* for SMFAs, with the remainder used for maintaining mosquito breeder populations until receipt of RBCs from the next donor.

In a broader context, if growth and differentiation of *P. falciparum* is reduced in RBCs that have been refrigerated for different durations, i.e., >1-week, simultaneous comparisons with cryo-preserved blood may help identify physico-chemical changes in blood that are “naturally” detrimental to parasite fitness. In fact, the adverse effects of prolonged storage of refrigerated RBCs in some cases has led the FDA to recommend identifying the true shelf-life of refrigerated RBCs and towards this end, clinical researchers have started using “omics” technologies to extensively catalog storage-related physico-chemical artefacts, some of which could indeed serve as suitable starting points for comparison [23, 25]. Along the same lines, cryo-preservation has the potential to offer a superior alternative to using older, refrigerated RBCs where “storage-related artefacts”, possibly compounded by inherent differences in RBCs between donors, may influence the interpretation of assays quantifying parasite fitness and/or reproducibility across labs, for instance, while comparing mutant strains or testing for drug susceptibility [14]. Finally, if differences between RBC donors are reflected in the mechanisms and consequences of pathogenesis for instance, cryo-preservation would provide an ideal tool for replicating the observations or allow testing other parasite strains [33].

In summary, we anticipate our approach to be compatible with a broad range of research questions involving the biology of gametocytogenesis as well as human-to-mosquito transmission, with significant potential for integrating into existing research pipelines in addition to identifying novel avenues for target identification and therapeutic intervention. Blocking this stage of the parasite’s life-cycle is predicted to be one of the last remaining hurdles to overcome before we can declare a malaria-free world [1, 2], but for which SMFAs will continue to serve as the penultimate reference.

## Materials and Methods

### Study design

The overall study design is described below, and in the supplement accompanying this manuscript (additional file 1). In general, each experimental block in the current study progressed in the following manner: 1) gametocytogenesis of *P. falciparum* NF54/CB132 was initiated in refrigerated (4°C) or cryo-preserved RBCs *in vitro*, 2) cultures were maintained for 14-16 days, and 3) fed to mosquitoes in SMFAs to assess the efficiency of gametogenesis and sporogony (additional file 2a).

To determine if cryo-preservation alters the ability of RBCs to support *P. falciparum* NF54 gametocytogenesis and mosquito infection, a nested/fully-crossed experimental design was implemented with two unique donors of RBCs serving as the basis of biological replication in two independent experimental blocks (additional file 2b, “nested design”) [34-36]. To account for any technical/biological variation arising from unknown factors independent of RBC storage method, each experimental block fulfilled three important criteria. First, mature gametocytes were cultured in refrigerated (4°C) or cryo-preserved RBCs from the same donor to ensure that any differences observed were not due to inherent differences between donors. Second, gametocyte flasks for comparing the two methods were initiated from the same asexual seed culture since parasite fitness can vary over generations, which in turn can introduce variation in induction of gametocytogenesis. Third, infectious cultures were offered to mosquitoes that were sorted from the same starting population and age post-emergence to minimize any variation in sporogony caused by inter-generational differences across mosquito cohorts. In summary therefore, each experimental block for the comparison between ≤1-week old refrigerated and cryo-preserved RBCs would consist of the following steps: 1) the same asexual seed culture was used to initiate gametocytogenesis of *P. falciparum* NF54 in ≤1-week old refrigerated (4°C) or cryo-preserved RBCs *in vitro*, 2) cultures maintained and monitored in parallel for 14-16 days, and 3) fed to mosquitoes from the same cohort to assess the efficiency of gametogenesis and sporogony (additional file 2). Lastly, it should be noted that due to practical constraints, that the comparison between ≤1-week old refrigerated and cryo-preserved RBCs were performed after the latter had been stored for 2-3 days.

In the second part of this study, we determined if duration of storage affects the ability of cryo-preserved RBCs to support transmission-competent gametocytes by assessing gametocytogenesis and infectivity to mosquitoes for parasites cultured in RBCs thawed after cryogenic storage for 4, 8 and 12 weeks. Biological replication for each storage period was again provided by RBC donors wherein 2-3 donors were tested independently for their ability to support infectious gametocytes after cryogenic storage for 4, 8 and/or 12 weeks. To fulfil the second and third criteria outlined above, we were unable to determine the effect of storage duration across all RBC donors, which resulted in a partially crossed experimental design (additional file 2b, “partially crossed design). In this part of the study, each experimental block was defined by the following steps: 1) the same asexual seed culture was used to initiate gametocytogenesis of *P. falciparum* NF54 in cryo-preserved RBCs from 2-3 donors at 4, 8 and/or 12 weeks of preservation with independent donors representing each storage period, 2) cultures maintained and monitored in parallel for 14-16 days, with 3) mature gametocytes from each flasks offered to mosquitoes from the same cohort and assessed for efficiency of gametogenesis and sporogony (additional file 2).

### Chemicals and consumables

All reagents and consumables described herein were purchased from Fisher Scientific (Hampton, NH) unless noted otherwise.

### Preparation of media components

Parasite culture media was prepared and stored as described previously [15], with minor modifications. Briefly, incomplete media was prepared by dissolving pre-made RPMI-1640 powdered media in distilled water before adding 2% sodium bicarbonate (w/v) and 0.005% hypoxanthine (w/v, Sigma-Aldrich, St. Louis, MO). The incomplete media was then filter-sterilized with 0.2 µm filters under vacuum before moving to storage at −20°C in aliquot sizes of 450 or 900 mls. Complete media was prepared just prior to use by adding 50 or 100mls of 10% A+ non-heat inactivated human serum (Valley Biomedical, Winchester, VA) to 450 or 900mls to thawed, incomplete media respectively. Media was dispensed into 45 ml aliquots in 50 ml conical-bottom tubes and the air-liquid interface sparged for 5-10 seconds with a micro-aerophilic gas mixture of 5% CO2, 5% O2 and 90% N2 (referred to herein as “tri-gas mixture”, Airgas LLC, Kennesaw, GA) prior to storage for 1 week. Since “quality” of serum is critical for culturing transmission competent parasites, serum samples were tested as described previously [15]. Serum from 14 individuals was pooled into pairs and tested for their ability to support gametocytogenesis as well as ex-flagellation of male gametocytes in case of *P. falciparum* isolate NF54 (unpublished observations). The same pool of serum donors was used to culture *P. falciparum* NF54 for the duration of this study.

### Cryo-preservation and thawing of RBCs

Cryo-preservation of RBCs was performed 3-4 days after collection as suggested elsewhere [27, 37]. Procedures for freezing and thawing were adapted from Sputtek [37] and Goheen et al [28] with minor modifications. Two-fold concentrated stocks of cryo-protectant were prepared in deionized water by warming a solution of 28% glycerol (v/v), 3% sorbitol (v/v from a 1M stock) and 0.65% (w/v) sodium chloride (NaCl) prior to sterilization with a 0.2 µm filter and storage at room temperature. Whole blood (∼500ml units, Valley Biomedical, Winchester, VA or Interstate Blood Bank, Memphis, TN) was dispensed in 45 ml aliquots and either stored refrigerated at 4°C until use (see next section) or allowed to warm to room temperature prior to cryo-preservation. To ensure reproducible recovery of viable RBCs following cryo-preservation, it is critical for the cryo-protectant and RBCs to be at the same temperature (e.g., room temperature), prior to use [37].

Aliquots of whole blood, no more than 3-4 days post-collection, were centrifuged at 1800xg for 10 minutes at room temperature at a low brake setting and plasma and white blood cell layers aspirated under vacuum before an equal volume of cryo-protectant solution (∼20-25mls) was added to the packed RBC pellet to achieve a final glycerol concentration of 14% (v/v) and a hematocrit of ∼50%. The packed RBCs were then equilibrated with the cryo-protectant for 15-20 minutes with gentle intermittent mixing. Glycerolized RBCs were then dispensed in 2 ml aliquots and snap-frozen by immersing in liquid nitrogen for 2-3 minutes before transferring to the vapor phase for long-term storage. Cryo-preserved RBCs were thawed just before use by incubating cryogenic vials in a dry bath for 5 minutes at 37°C before transferring the contents into a 15 or 50 ml conical bottom tube prior to de-glycerolization. Unless stated otherwise, all solutions for de-glycerolization were prepared in deionized water and sterilized with 0.2µm filters before storage at room temperature. De-glycerolization was initiated with the addition of successive gradients of sodium chloride concentrations starting with 0.4mls (0.2 volumes) of 12% NaCl (w/v in distilled water) added dropwise under gentle agitation (∼1 minute). The mixture was then incubated at room temperature for 5 minutes with gentle intermittent mixing before dropwise addition of 10mls (5 volumes) of sterile 1.6% NaCl (w/v in distilled water). The de-glycerolized RBC suspension was centrifuged at 1000xg for 3 minutes at 20-23°C and low brakes and 20mls (10 volumes) of a salt-dextrose solution (0.9% NaCl + 0.2% dextrose, w/v in distilled water) added dropwise to the packed RBC pellet under gentle agitation. If hemolysis was still observed in the supernatant, the packed RBCs were washed once more with the same volume of salt-dextrose solution. Finally, the thawed RBCs were resuspended in complete media before use. Recoveries of ∼0.8-0.9mls of packed RBCs should be considered routine.

### Preparation of refrigerated RBCs

On the day of use, but ≤1-week post-collection, a 45 ml aliquot of whole blood was retrieved from the refrigerator and allowed to warm to room temperature before centrifugation as described in the preceding section. The plasma and white blood cell layers were aspirated before the addition of an equal volume of incomplete media (∼20-25mls), also at room temperature. RBCs were gently suspended in incomplete media before centrifugation for 3 minutes at 1000xg and low brake setting. Packed RBCs were washed two more times before resuspension in an equal volume of complete media to achieve a final hematocrit of ∼50%.

### Cultures of asexual stage *P. falciparum*

*P. falciparum* isolate NF54 was obtained from BEI Resources, NIAID, NIH (“*Plasmodium falciparum*, Strain NF54 (Patient Line E), product number MRA-1000, contributed by Megan G. Dowler”). Asexual feeder cultures were routinely sub-cultured in T25 or T75 flasks in a volume of 5 or 15mls of complete media respectively, with final hematocrit of 5% and ring state parasitemia ranging from 0.1-1%. The same volume of media was replaced every 24 hours and then sparged gently at the air-liquid interface with the tri-gas mixture for 15-20 seconds. Parasitemia was monitored every 1-2 days by transferring 50-100 µl of parasite culture to a 0.6ml tube, centrifuged at 1800xg for 1 minute and supernatants aspirated until final culture volume was ∼15-20 µl. The RBC pellet was re-suspended with repeated pipetting before preparing smears with 2 µl of the concentrate on glass slides. Slides were dried on a slide-warmer for 20-30 seconds and then fixed by submerging in 100% Methanol for 10 seconds prior to Giemsa staining. Concentrated Giemsa stain (Sigma-Aldrich, St. Louis, MO) was filtered in aliquot sizes of 30-35mls with a Whatman no. 1 filter paper and stored at room temperature. Just before use, the filtered stain was diluted to a ratio of 1:20 (v/v) in phosphate-buffered saline (pH 7.2) and fixed slides immersed in the stain in glass Coplin-jars for 10-15 minutes at room temperature. Stained sides were washed under running tap-water for 5-10 seconds with the smeared side facing downwards before being air-dried for microscopy. Smears were mounted directly in immersion oil (Cargille Labs, Cedar Grove, NJ) and examined at 1000x magnification with a Leica DM2500 upright microscope (Leica Microsystems, Buffalo Grove, IL). Parasitemia was estimated from ∼1000 RBCs and staged into early trophozoites (referred to herein as rings), late trophozoites and schizonts following guidelines described previously [13, 38]. Ring-stage parasitemia expressed as a proportion of RBCs counted was used to prepare flasks for routine sub-culture and/or gametocytogenesis [15].

### Cultures of the sexual stages (gametocytes) of *P. falciparum*

Flasks destined for gametocytogenesis were only initiated when ring-stage parasitemia in the asexual seed flasks above showed exponential increase in growth rates (>6-8 fold) relative to the previous sampling point. Flasks for gametocytogenesis were prepared essentially as described above. Briefly, freshly thawed or refrigerated RBCs resuspended in pre-warmed (37°C) complete media at 5% hematocrit (e.g., 1mls naïve RBCs in 20mls complete media) were seeded with asexual cultures to achieve a final ring-stage density of 0.6-1% in the gametocyte flasks and returned to an incubator maintained at 37°C. Media in flasks was replaced every 24 hours with 15mls of fresh pre-warmed (37°C) complete media and sparged with tri-gas mixture for 10-30 seconds. All manipulations outside the incubator were performed by placing the flask on a slide-warming platform maintained at 38°C (MedSupply Partners, Atlanta, GA). Gametocyte flasks were maintained for 14-16 days and parasitemia monitored with Giemsa staining at 1, 4, 8, 12 and 14 days post-infection as suggested previously [15]. Asexual stages were staged as described above and gametocytes were further classified into stages II, III, IV and V male or V female gametocytes using guidelines established previously [13, 38] (additional file 4a and S4b). Stage I gametocytes were excluded from the counts as they are virtually indistinguishable morphologically from late trophozoites, especially in asynchronous cultures [39]. Finally, maturation status of the culture was assessed from 12 days post-infection by quantifying ex-flagellation of male stage V gametocytes in a Neubauer hemocytometer chamber as suggested previously [15]. Briefly, after adding fresh media, 10 µl of culture was pipetted directly into the chambers of a hemocytometer and incubated at 21-24°C for 15-20 minutes before quantification (additional file 4c). Imaging was performed at 400x magnification on a Leica DM2500 equipped with differential interference contrast (DIC) optics. Cultures were deemed infectious only when ex-flagellation was observed and offered to mosquitoes within 24-48 hours.

*P. falciparum* Cambodian isolate CB132 (kind gift of Prof. Dennis E Kyle, University of Georgia, Athens, GA, USA) [30] was cultured under similar conditions to NF54 with the exception that flasks destined for gametocytogenesis were seeded at a density of 0.5%.

### Mosquito husbandry

*An. stephensi* mosquitoes were maintained in a level 2 Arthropod Containment Laboratory at the University of Georgia, which were initiated from eggs kindly provided by the Walter Reed Army Institute of Research ca. 2015 is a wild-type strain referred to as “Strain Indian” [10, 40]. Colonies were housed in a dedicated walk-in environmental chamber (chamber (Percival Scientific, Perry, IA) at 27°C + 0.5°C, 80% +5% relative humidity, and under a 12 hr light: 12 hr dark photo-period schedule. Adult mosquitoes were maintained on 5% dextrose and provided whole human blood offered in glass-jacketed feeders (Chemglass Life Sciences, Vineland, NJ) through parafilm membrane maintained at 37°C to support egg production. Husbandry procedures were established according to guidelines as suggested elsewhere [32] with minor modifications. Briefly, eggs were rinsed twice with 1% house-hold bleach (v/v, final concentration of 0.06% sodium hypochlorite) before surface-sterilization for 1 minute in the same solution at room temperature. Bleached eggs were washed with 4-5 changes of deionized water and transferred to clear plastic trays (34.6cm L × 21.0cm W × 12.4cm H) containing 500 ml of deionized water and 2 medium pellets of Hikari Cichlid Gold fish food (HikariUSA, Hayward, CA) and allowed to hatch for 48 hours. Hatched L1 larvae were dispensed into clear plastic trays (34.6cm L × 21.0cm W × 12.4cm H) at a density of 300 larvae/1000 ml water and provided the same diet until pupation. The feeding regime consisted of 2 medium pellets provided on the day of dispensing (day 0) followed by the provision of a further 2, 4, 4 and 4 medium pellets on days 4, 7, 8 and 9 respectively. This regime allows >85% larval survival and >90% pupation within 11 days with a sex ratio of 1:1 adult males and females (unpublished observations).

### Mosquito infections

All steps outlined below were performed with pre-warmed (38°C) equipment, including tubes, pipettes, pipette tips and centrifuge rotor buckets. Cultures deemed infectious were concentrated by centrifugation at 1800xg for 1-2 minutes at room temperature in pre-weighed 15 or 50 ml conical bottom centrifuge tubes. Supernatants were aspirated under vacuum and the weight/volume of the concentrate estimated by weighing the tube and subtracting the value of the pre-weighed empty tube. 3-6 volumes of a freshly washed and pre-warmed mixture of RBCs resuspended in freshly thawed serum (30% hematocrit) was added to the infected RBC pellet to achieve a final hematocrit of 45-50% and mature gametocytemia of ∼0.1%. The volumes of naïve RBCs and serum mixture to be added was informed by the densities recorded in the gametocyte flasks on the day of infection. The mixture of naïve and parasitized RBCs was then fed immediately to 60-80 (3-7 day old) female *An. stephensi* mosquitoes for 15-20 minutes. To facilitate high blood feeding rates, mosquitoes were previously starved for 16-24 hours in dedicated environmental chambers programmed to fluctuate a total of 9°C around a daily mean of 24°C with 80% ±5% relative humidity and a 12h light:dark photo-period schedule (Percival Scientific, Perry, IA) as described previously [10]. After the blood-feeds, feeding status was qualitatively ascertained and engorged mosquitoes selected for by extending the starvation period for a further 24-48 hours to eliminate non-blood-fed mosquitoes [41].

To quantify parasite densities in the infectious feed, Giemsa stained smears were prepared from a 2 µl aliquot of the infectious blood meal offered to mosquitoes, as described above. Prevalence and abundance of parasites in the midguts of mosquitoes was assessed at 9-13 days post-infection by dissecting midguts and counting oocysts at 400x magnification with a Leica DM2500 under DIC optics (additional file 4d). Sporozoite prevalence was assessed at 17-21 days post-infection by dissecting salivary glands into 5 µl of PBS. Glands were ruptured by overlaying a 22mm^2^ coverslip and presence/absence of sporozoites was recorded for each mosquito at either 100x or 400x magnification with the same microscope (additional file 4e).

### Statistical modeling and data analyses

All data analyses were performed in RStudio (version 1.1.423) [42] running the statistical software package R (version 3.5) [43]. Due to the experimental design (see above and figures S1 and S2), generalized linear mixed-effects models (GLMMs) were used for all statistical analyses, as suggested previously [34-36]. Choice of GLMM family and corresponding link function were based on five dependent variables modeled-1) rates of mature gametocytemia *in vitro*, 2) oocyst prevalence (proportion of mosquitoes with oocysts on the midgut), 3) oocyst abundance (total number of oocysts per midgut regardless of infection status), 4) oocyst intensity (number of oocysts per midgut from infected midguts only), and 5) sporozoite prevalence (proportion of mosquitoes with sporozoites in the salivary glands). For all the analyses, storage method (refrigerated vs. cryo-preserved RBCs) and duration (4, 8, and 12 weeks post cryo-preservation) were specified as independent variables, with storage method and duration classified as a categorical and continuous predictor, respectively. Storage duration was centered and scaled over the grand mean. For analyzing temporal patterns of gametocytogenesis *in vitro*, GLMMs with a binomial distribution (“logit” link) were performed with the total count of RBCs infected with male and female stage V gametocytes (i.e., total mature gametocytes) expressed as a proportion of the sum of RBCs (infected and uninfected) counted from each sample as the dependent variable. Oocyst and sporozoite prevalence in the midguts and salivary glands of mosquitoes, respectively, were modeled as the probability of mosquitoes being infected and infectious as dependent variables in GLMMs specified with a binomial distribution (“logit” link function). For analyzing oocyst abundance and intensity, GLMMs with negative binomial distributions (“log” link) were utilized. All statistical analyses were performed using the package “lme4” unless stated otherwise [44]. For the *in vitro* analysis, the intercepts of the relationships between the rates of mature gametocytemia and the predictor variables was allowed to vary between RBC donors as well as over sampling period (days post-infection) between experimental blocks and flasks nested within each block. The same random effect structure was specified for analyzing parasite fitness in the mosquitoes except infections/SMFAs were nested within blocks *in lieu* of flasks (additional file 2b). Tabulation of model outputs, along with tests for overdispersion were performed with the “sjstats” package. Where applicable, post-hoc comparisons of storage method and duration were performed using Tukey contrasts of the estimated marginal means derived from the models above, as suggested in the “emmeans” package [45].

Finally, cross-validation of the models were performed by 1) testing fit after truncating the number of mosquitoes sampled across each infection and 3) removing experimental block(s) and/or flasks/infections within each block. Where applicable, random effects were checked for normal distribution patterns and the final output tables of statistical modeling prepared with the “sjPlot” package and “sjstats” packages [46, 47]. All figures were prepared using the “ggplot2” package [48].

## List of abbreviations

DIC: differential interference contrast
GLMMs: generalized linear mixed-effects models
IRR: incidence rates ratios
OR: odds ratios
SMFAs: small membrane feeding assays

---

**Additional file 1.**
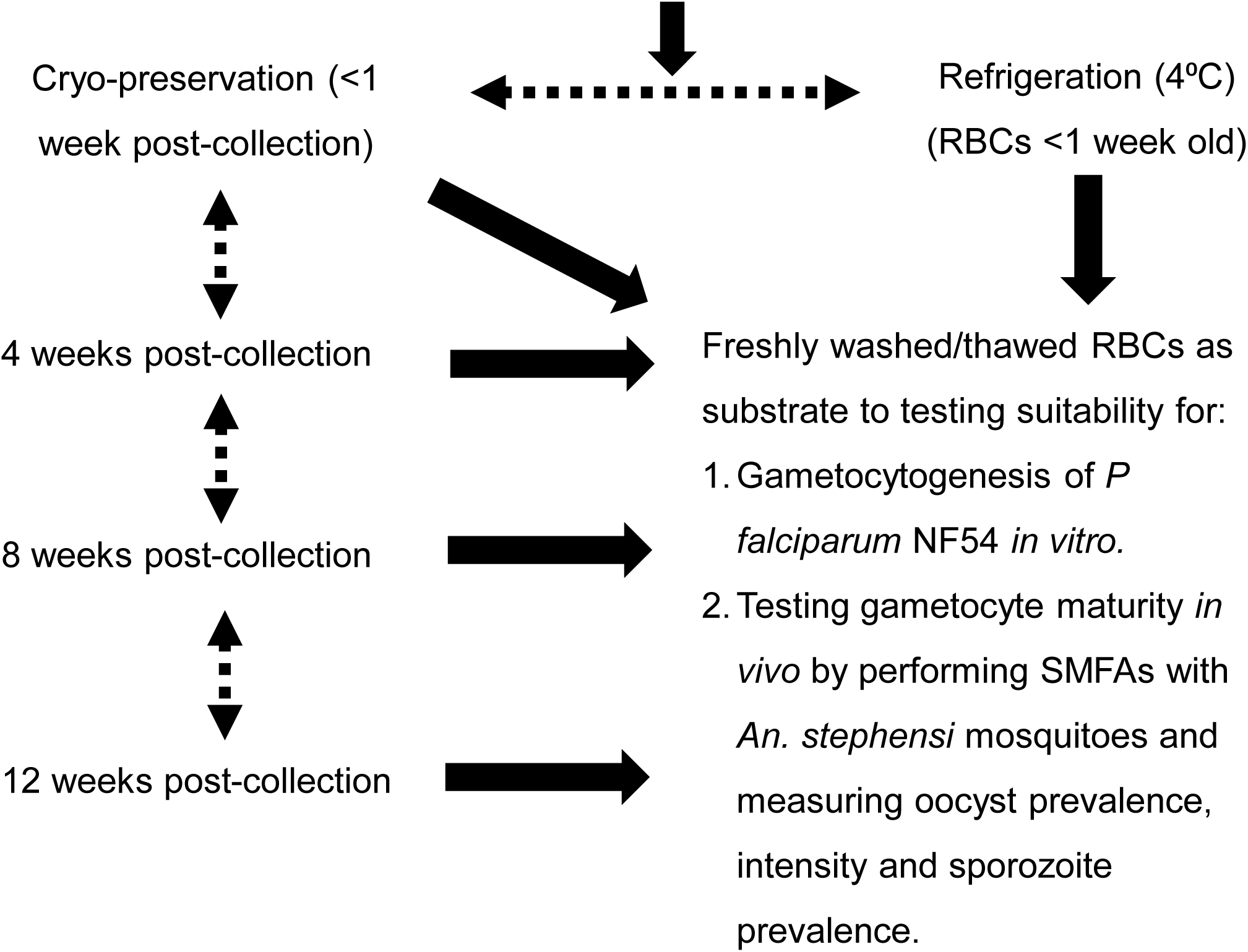
File format: “pdf”. Title of data: A schematic of the overall study design. Description of data: RBCs 3-4 days old post-collection were either refrigerated at 4°C or cryo-preserved in the gaseous phase of liquid nitrogen. Aliquots of RBCs were thawed at 1, 4, 8 and 12 weeks and assessed for their ability to support 1) gametocytogenesis of *P. falciparum* NF54 *in vitro* and 2) gametogenesis *in vivo* relative to refrigerated RBCs which served as the reference (dashed arrows). Black continuous arrows indicate procedures that were common to all treatments.

**Additional file 2:**
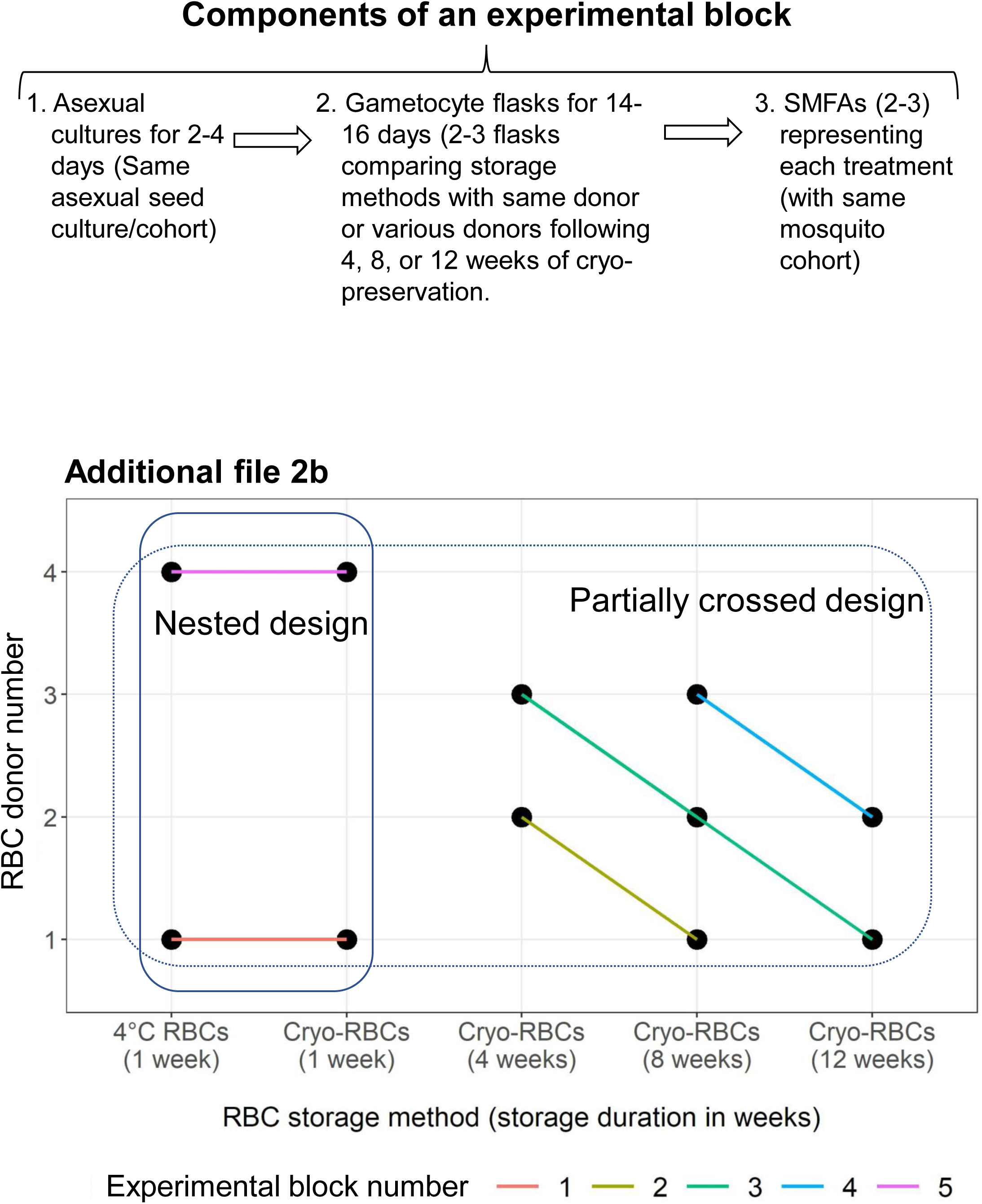
File format: “pdf”. Title of data: A detailed schematic of the experimental design describing biological replication and testing regime. Description of data: a.) The sequence of steps in an experimental block with design objectives in parentheses. See main text for details. b.) Testing regime to demonstrate how RBCs from 4 independent donors were used to compare storage method (refrigeration vs. cryo-preservation) and duration of storage. The colored lines connecting each data point indicate which donors were tested within an experimental block/unit. While the comparison between storage methods was performed for donors 1 and 4 in a nested experimental design, the comparisons for storage duration encompassed a partially crossed design with at least 2-3 donors tested at 4, 8 and 12 weeks respectively with repeated sampling of 3 of the 4 total donors at various periods following cryo-preservation. For instance, RBCs from donor 3 was tested following 4 and 8 weeks of cryo-preservation in experimental blocks 3 and 4 respectively. Additionally, in block 3 (green lines), donors 1 and 2 were also tested after cryo-preservation for 12 and 8 weeks respectively along with donor 3 whose RBCs had been stored for 4 weeks by then.

**Additional file 3:**
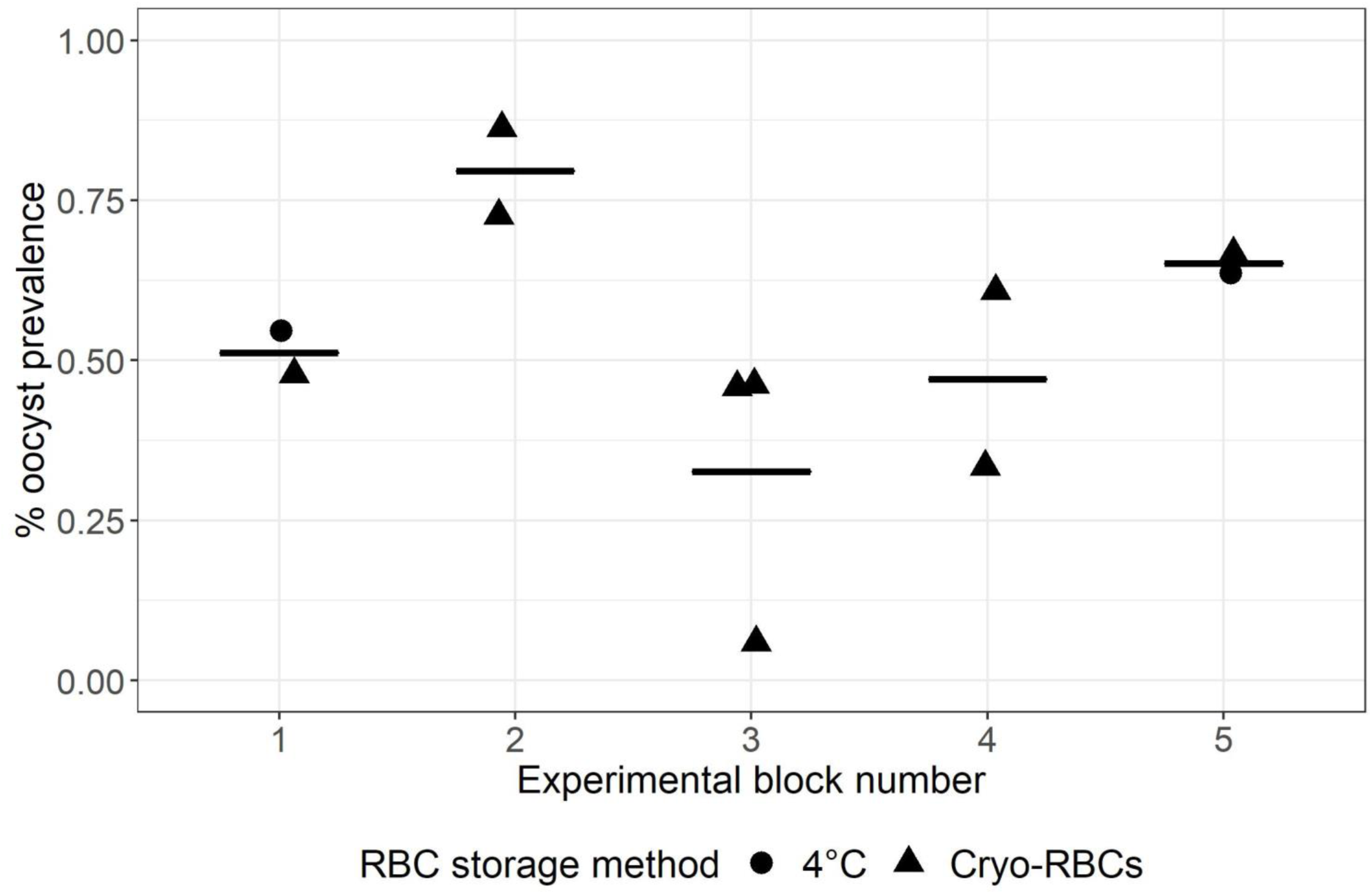
File format: “pdf”. Title of data: Variation between experimental blocks in the prevalence of *P. falciparum* NF54 oocysts in midguts of *An. stephensi*. Description of data: Each data point within an experimental block represents an individual SMFA performed within the same block as depicted in Additional file 2. Horizontal bars represent mean prevalence within each block.

**Additional file 4:**
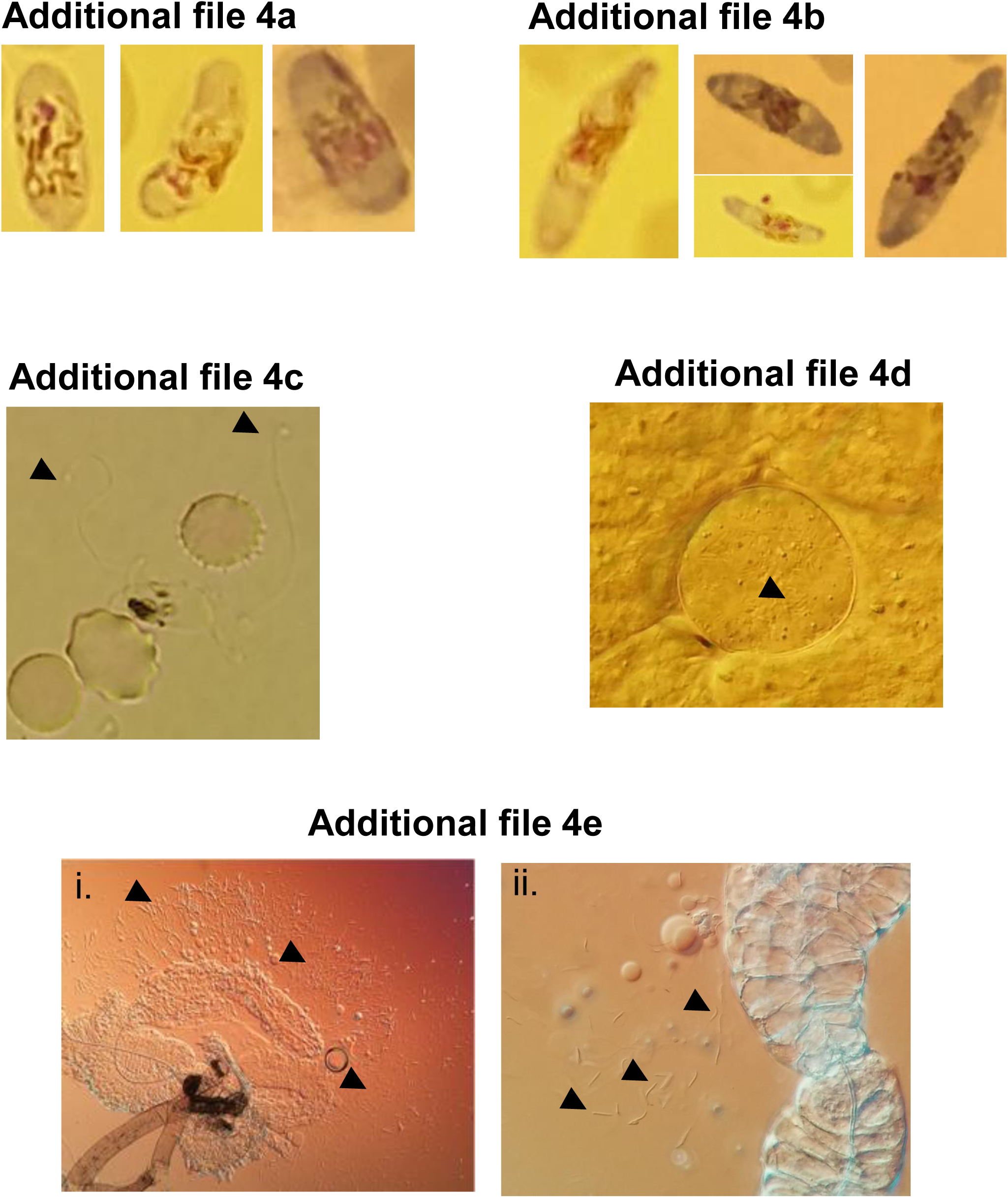
File format: “pdf”. Title of data: Representative images used to classify the various stages of *P. falciparum* NF54 during this study. Description of data: a.) Giemsa-stained images of male gametocytes (1000x, oil immersion, brightfield), b.) Giemsa-stained images of female gametocytes (100x, oil immersion, brightfield), c.) Ex-flagellation of gametocytes *in vitro* with arrowheads depicting flagella (400x DIC, unstained), d.) An oocyst with enclosed sporozoites (arrowheads, 400x, DIC, unstained), e.) ruptured salivary glands with freed sporozoites (arrowheads) at 100x (i.) and 400x (ii) (DIC, unstained). Images were captured with an LG G3 or Samsung Galaxy S7 smartphone using default settings (“Auto”) while attached to the eyepieces on the microscope with a custom-designed apparatus. Except panel e, all images were digitally magnified up to 4x for presentation purposes.

**Additional file 5:**
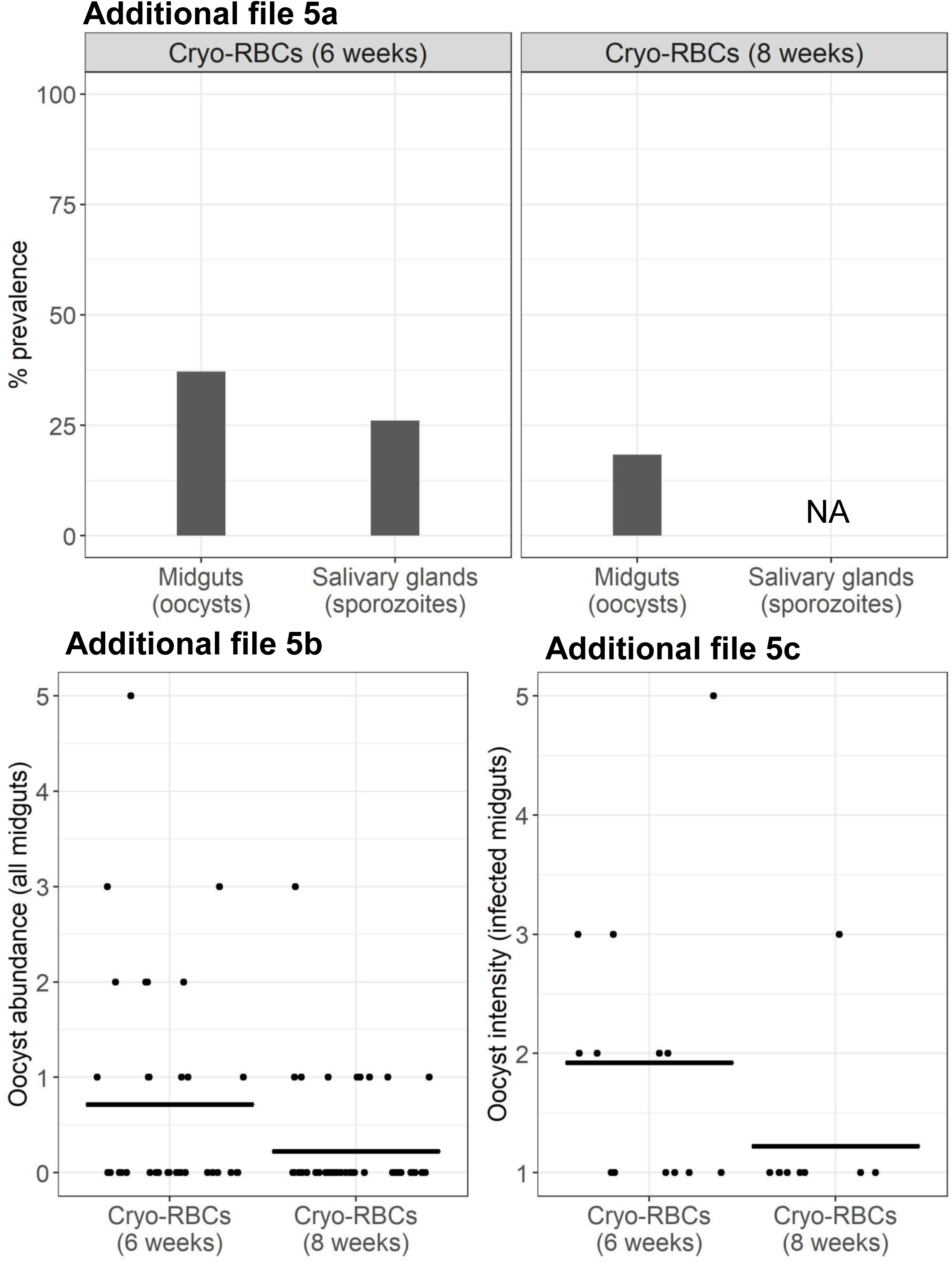
File format: “pdf”. Title of data: Cryo-preserved RBCs support SMFAs with a Cambodian isolate of *P. falciparum*. Description of data: a) Oocyst and sporozoite prevalence, b) oocyst abundance and c) intensity of *P. falciparum* CB132 in female *An. stephensi* infected with mature gametocytes of *P. falciparum* CB132 cultured in RBCs from donor 4 (Additional file 2) thawed following cryo-preservation for 6 (left panel) or 8 weeks (right panel). Female 3 to 5-day old *An. stephensi* mosquitoes were provided an infectious blood-meal spiked with ∼0.6% mature gametocytes and oocyst prevalence and abundance recorded at 12 days post-infection and sporozoite prevalence at 17 days post-infection. Horizontal bars represent group means with each data point representing oocyst counts from an individual mosquito midgut. For visualization purposes, counts were jittered horizontally to 40% but not vertically to maintain alignment with the gradient on the y-axis. NA=not available.

